# Measurement of *Klebsiella* Intestinal Colonization Density to Assess Infection Risk

**DOI:** 10.1101/2021.02.16.431551

**Authors:** Yuang Sun, Alieysa Patel, John SantaLucia, Emily Roberts, Lili Zhao, Keith S. Kaye, Krishna Rao, Michael A. Bachman

## Abstract

**Background:** *Klebsiella pneumoniae* and closely related species *K. variicola* and *K. quasipneumoniae* are common causes of healthcare-associated infections, and patients frequently become infected with their intestinal colonizing strain. To assess the association between *Klebsiella* colonization density and subsequent infections, a case-control study was performed.

**Methods:** A multiplex qPCR assay was developed and validated to quantify *Klebsiella (K. pneumoniae, K. variicola, and K. quasipneumoniae* combined) relative to total bacterial DNA copies in rectal swabs. Cases of *Klebsiella* infection were identified based on clinical definitions and having a clinical culture isolate and preceding or co-incident colonization isolate with the same *wzi* capsular sequence type. Controls were colonized patients without subsequent infection and were matched 2:1 to cases based on age, sex, and rectal swab collection date. Quantitative PCR (qPCR) from rectal swab samples was used to measure the association between relative abundance (RA) of *Klebsiella* and subsequent infections.

**Results:** *Klebsiella* RA by qPCR highly correlated with 16S sequencing (ρ=0.79; *P* <.001). The median *Klebsiella* RA in the study group was 2.6% (interquartile range (IQR) 0.1-22.5, n=238), and was higher in cases (15.7%, IQR 0.93-52.6%, n=83) than controls (1.01%, IQR 0.02-12.8%; n=155; *P* <0.0001). After adjusting for multiple clinical covariates using inverse probability of treatment weighting, subjects with a *Klebsiella* RA > 22% had a 2.87-fold (1.64-5.03, *P* =0.0003) increased odds of infection compared to those with lower colonization density levels.

**Conclusions:** Measurement of colonization density by qPCR could represent a novel approach to identify hospitalized patients at risk for *Klebsiella* infection.

**Importance:** Colonization by bacterial pathogens often precedes infection, and offers a window of opportunity to prevent these infections. *Klebsiella* colonization is significantly and reproducibly associated with subsequent infection, however factors that enhance or mitigate this risk in individual patients are unclear. This study developed an assay to measure the density of *Klebsiella* colonization, relative to total fecal bacteria, in rectal swabs from hospitalized patients. Applying this assay to 238 colonized patients, high *Klebsiella* density defined as >22% of total bacteria, was significantly associated with subsequent infection. Based on widely available polymerase chain reaction (PCR) technology, this type of assay could be deployed in clinical laboratories to identify patients at increased risk of *Klebsiella* infections. As novel therapeutics are developed to eliminate pathogens from the gut microbiome, a rapid *Klebsiella* colonization density assay could identify patients who would benefit from this type of infection prevention interventions.

## Introduction

*Klebsiella pneumoniae* is a leading cause of healthcare-associated infections (HAIs)(1). Recent studies have shown that *Klebsiella variicola* and *Klebsiella quasipneumoniae* are closely related to, yet distinct species from, *K. pneumoniae* and cause indistinguishable infections (2, 3). These species are part of the *K. pneumoniae* complex that together pose a serious public health threat.

*Klebsiella* commonly colonizes hospitalized patients and can cause bacteremia, pneumonia, and urinary tract infections (UTIs). Prior studies show that *Klebsiella* colonization is significantly associated with subsequent infections and 80% of infections in colonized patients are caused by an intestinal colonizing strain (4, 5). Increased colonization density may increase the risk of subsequent infection. For example, intestinal domination (defined as at least 30% relative colonization density) of *Proteobacteria* was associated with subsequent Gram-negative bacteremia in patients undergoing allogeneic hematopoietic stem cell transplantation and relative and absolute abundance of *Enterobacterales* associate interactively with infection in intensive care patients (6, 7). In long term acute care patients, relative abundance of carbapenem-resistant *K. pneumoniae* above 22% was a risk factor for bacteremia (8). Similarly, increased relative abundance of *Escherichia* and *Enterococcus* in the gut are risk factors for corresponding bacteriuria or UTI in kidney transplant patients (9). These studies indicate that in many cases colonization is a necessary intermediate step before infection.

Understanding the association between *Klebsiella* colonization and subsequent infections could provide opportunities for identification of high-risk patients, intervention, and ultimately prevention of infection. Additionally, little is known about the association between *Klebsiella* gut colonization density and specific infection types, such as bacteremia, pneumonia, and UTI. Measuring *Klebsiella* gut density and assessing gut density as a risk factor for various infections may also shed light on the mechanisms of dissemination from the colonized gut to various infection sites. However, the lack of a rapid and reliable assay to quantify *Klebsiella* relative abundance in the gut has been a hindrance to both research and potential clinical implementation. Here we report a qPCR-based assay that can quickly and accurately quantify *Klebsiella* from rectal swab specimens. We employed this assay in a case-control study to assess *Klebsiella* rectal relative abundance as a risk factor for bacteremia, pneumonia or UTI in colonized patients and found a significant association after adjusting for clinical variables.

## Results

### *In silico* Analysis

To design a quantitative PCR (qPCR) assay for the *K. pneumoniae* complex, 31 *K. pneumoniae*, *K. quasipeumoniae*, and *K. variicola* strains with complete genomes were selected as “inclusivity” for *in silico* analysis (**Supplemental Dataset 1**). Additionally, 8 *K. oxytoca* and *Raoultella* strains were selected to represent “near-neighbor exclusivity”, and the human genome and common members of the gut microbiome were used as background sequences that should not be detected by the assay. PanelPlex™ *in silico* analysis was performed and the *ybiL* gene was identified as an optimal target for the assay (**Table 1**). An overall performance score, based on primer and probe thermodynamic stabilities with their targets, as well as any off-target bindings, were computed to constitute overall performance scores for each of 7 assay designs. The *ybiL* assay design 1 (overall score 99.9) has a predicted probe binding score of 99.5% with all 31 strains in the “inclusivity” set. In regard to the primers, all 31 predicted binding scores of the forward primer (hereafter *ybiL*-F) are above 50.0%, with 29 above 86.0%. Twenty-four predicted binding scores of the reverse primer (hereafter *ybiL*-R) are above 95.0% while 6 are close to 50.0% and 1 below 50.0%. Additionally, *ybiL* assay design 1 was predicted to have no amplifications with any background genomes. Although its primer bindings have variations, its probe binding scores are uniformly excellent. Therefore, *ybiL* assay design 1 (hereafter *ybiL* assay) was chosen for further validation. To assess the relative abundance of the *K. pneumoniae* complex, the *ybiL* assay and a previously described panbacterial qPCR assay targeting the 23S rRNA gene (sepsis manuscript) were combined to construct a multiplex qPCR assay (hereafter Kp qPCR assay). Overall, the *ybiL* assay has good coverage of the *K. pneumoniae* complex and the Kp qPCR assay provides a possible solution to quantify the *K. pneumoniae* complex in clinical specimens.

**Table 1.**
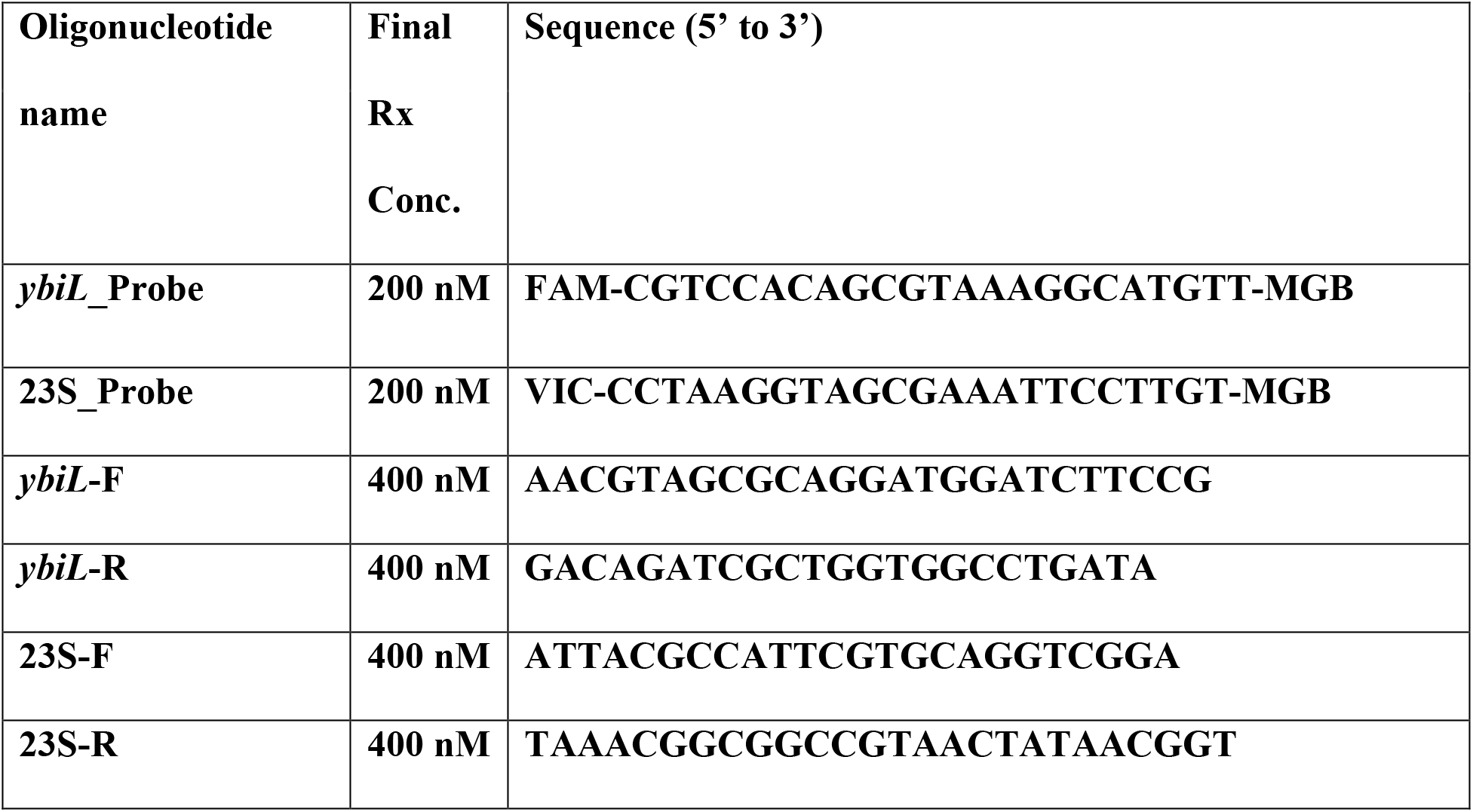
Primers and probes used in the study.

**Table 2.**
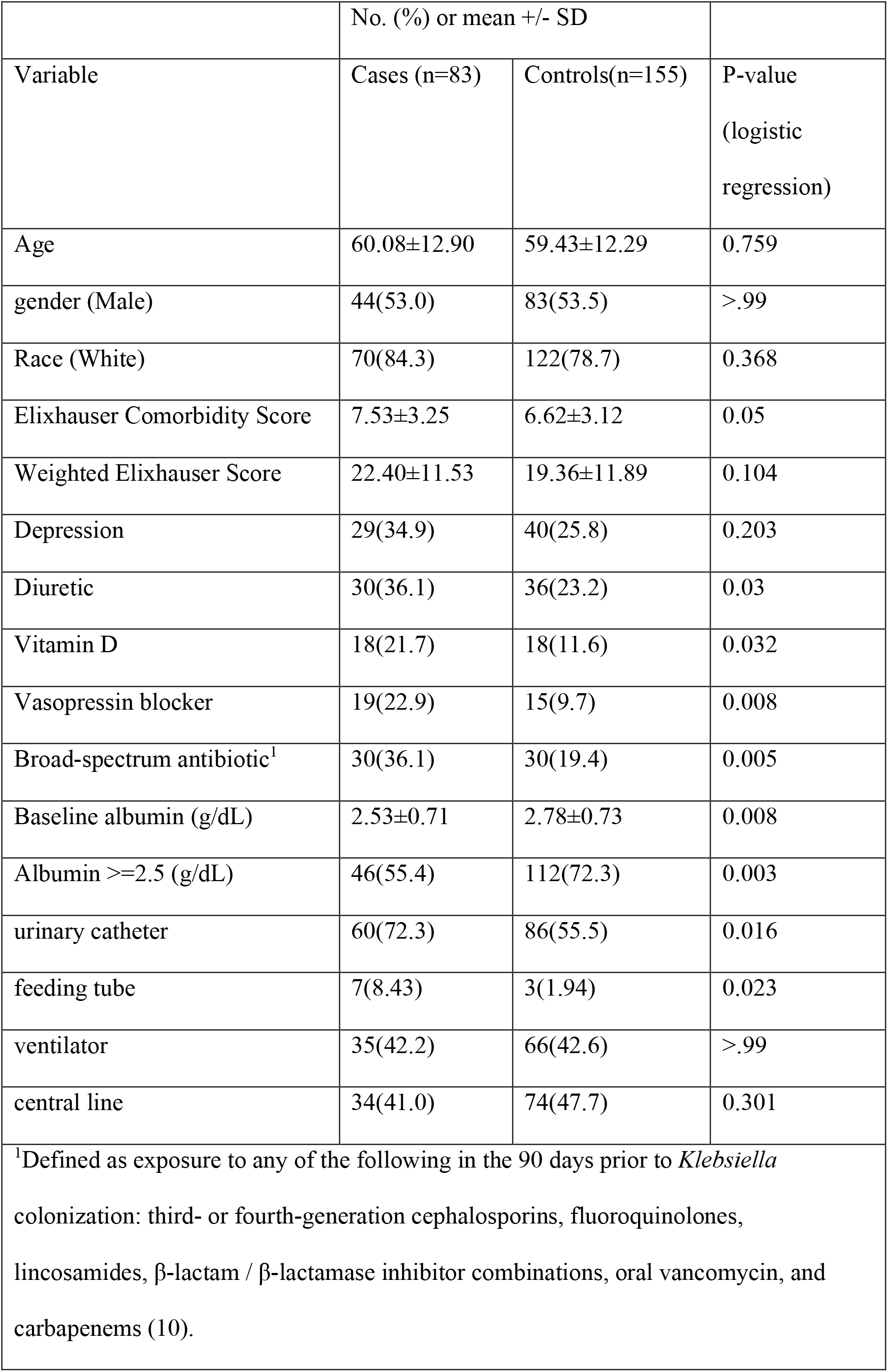
Patient Demographics

### *K. pneumoniae* Complex Diversity Panel

Eleven isolates with polymorphisms at sites of *ybiL* primer and probe binding were picked and grown overnight in LB broth. They were re-suspended in Amies media (BD ESwab™) and normalized based on colony forming units (CFU) for DNA extraction. The lab strain ATCC 43816 KPPR1, which contains a single mismatch identical to Kp8399, was set as reference and delta-delta-C_T_ (ddC_T_) method was used to calculate the abundance of *Klebsiella* relative to KPPR1 (set as 100%) by qPCR. (**Figure 1)** Of the 11 isolates, 9 are within the range of 88 to >99% relative to KPPR1. Although they share the same polymorphism, the abundance calculation of Kp2950 was 72% relative to KPPR1 whereas Kp6966 was 88%. This suggests that technical imprecision may be greater than systematic errors caused by polymorphisms. Taken together, the Kp qPCR assay should have accurate and consistent performance with most clinical isolates despite the existence of binding variations.

**Figure 1.**
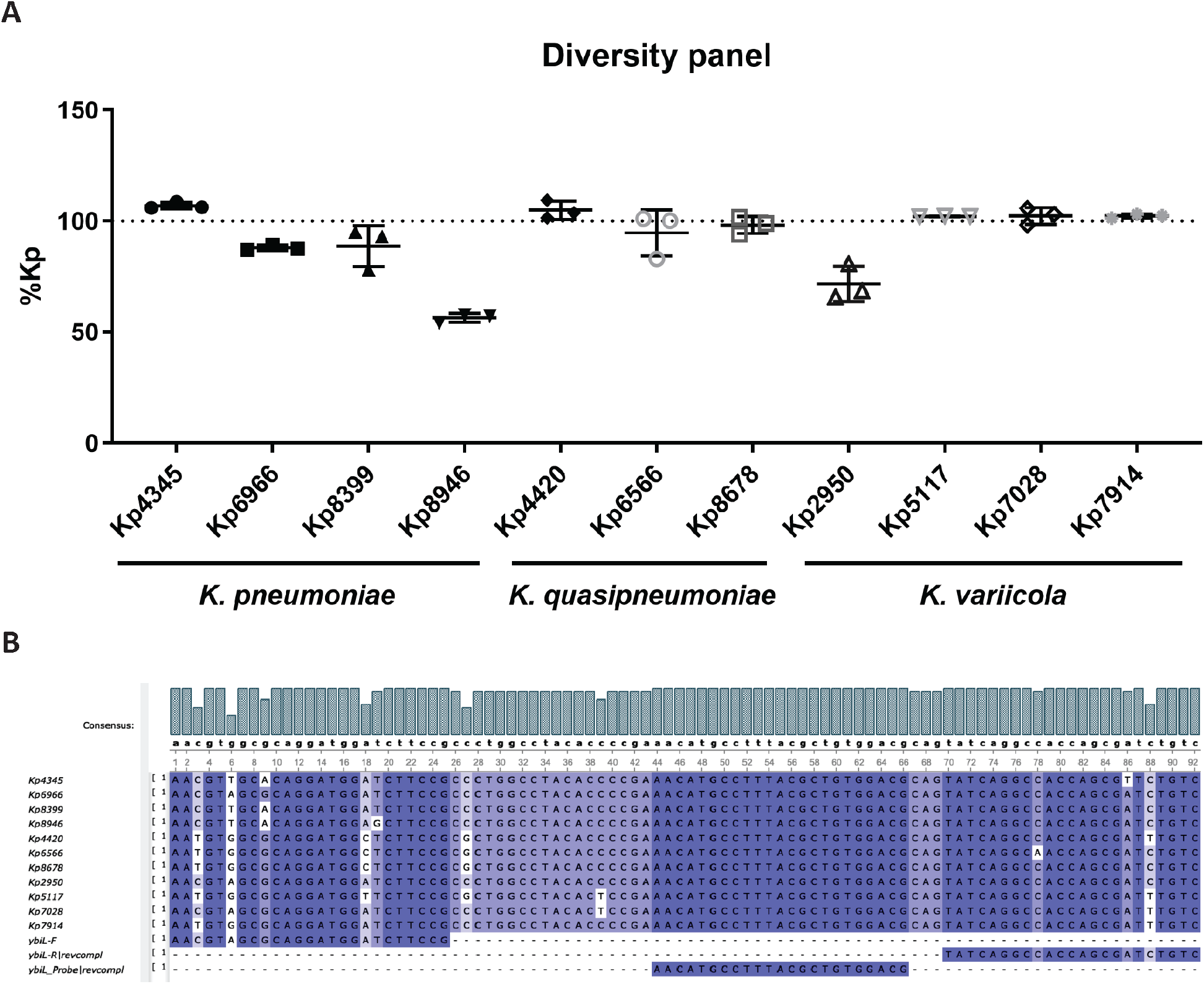
The Kp qPCR assay accurately quantifies *K. pneumoniae*, *K. variicola* and *K. quasipneumoniae*. Eleven *Klebsiella* clinical isolates were extracted and amplified by Kp qPCR assay, each with 3 technical replicates. A. Quantification of each isolate relative to KPPR1 set as 100%. B. The alignment of the amplicons of the 11 isolates with Kp qPCR assay primers and probe.

**Figure 2:**
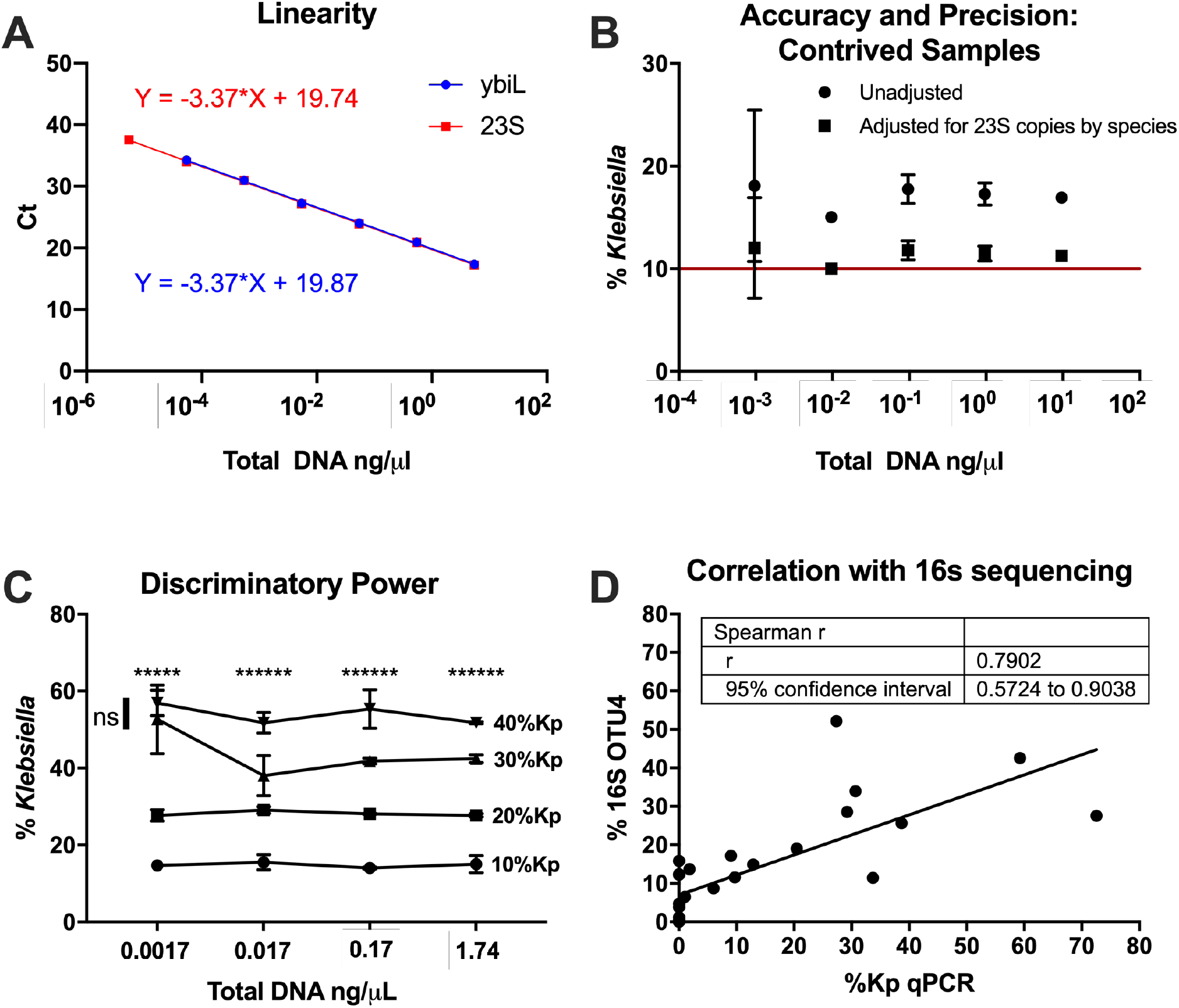
The Kp qPCR assay has the accuracy, precision and linearity to distinguish small differences in *Klebsiella* relative abundance. Linearity was assessed with serial dilutions of KPPR1 genomic DNA (n=3 technical replicates). Linear regression of Ct values from the *ybiL* assay (blue) and 23S (red) is shown (A). Precision and accuracy was assessed with serial dilutions of a mixture of 89% *Bacteroides ovatus*, 10% *K. pneumoniae* KPPR1, and 1% *Serratia marcescens* genomic DNA (3 technical replicates). Mean and SD of both direct and adjusted quantifications, after consideration of 23S gene copy number variations, are shown B). The ability to discern relative differences is shown using serial dilutions of mixtures of KPPR1 and *Escherichia coli* CFT073 (3 technical replicates). For each dilution, one-way ANOVA was performed (*P*<0.0001 for all) and Tukey’s post test was performed (* for each comparison out of six with *P*<0.05) (C). Accuracy was compared to 16S rRNA sequencing using OTU 4 that contains *Klebsiella*. The correlation between *Klebsiella* relative abundance by Kp qPCR and OTU4 of 16S rRNA sequencing analysis that contains *Klebsiella* was measured by Spearman’s rank correlation coefficient on 26 rectal swab samples (D).

### Specificity

The Kp qPCR assay was designed to quantify *K. pneumoniae*, *K. quasipeumoniae*, and *K. variicola* but not other *Klebsiella* species or other common bacteria in the gut microbiota. To validate its specificity, *K. aerogenes, K. pneumoniae subsp. ozaenae, K. oxytoca, Raoultella planticola*, *Raoultella ornithinolytica*, *Escherichia coli,* and *Pseudomonas aeruginosa* were tested by the Kp qPCR assay. (**Supplemental Table 1**) *K. pneumoniae* KPPR1 was used as positive control. Only KPPR1 and *K. pneumoniae subsp. ozaenae* were amplified by both *ybiL* assay and 23S assay, whereas *K. aerogenes, K. oxytoca, Raoultella planticola, Raoultella ornithinolytica, Escherichia coli, and Pseudomonas aeruginosa* strains were only amplified by 23S assay but not *ybiL* assay, demonstrating that *ybiL* assay specifically amplified the designated targets but not its near-neighbors or background sequences.

### Linearity

To assess Kp qPCR assay’s linearity, KPPR1 was grown in LB broth overnight and re-suspended in Amies media. A serial 10-fold dilution was made in triplicate and enumerated for CFU counts. The CFU counts were close to a theoretical 10-fold dilution, as the slope was 0.9889 and R square 0.9999. The slopes of *ybiL* and 23S assay were both 3.37 and R squares were both 0.9994, demonstrating that both assays had good linearity and efficiency. (**Figure 3**)

**Figure 3.**
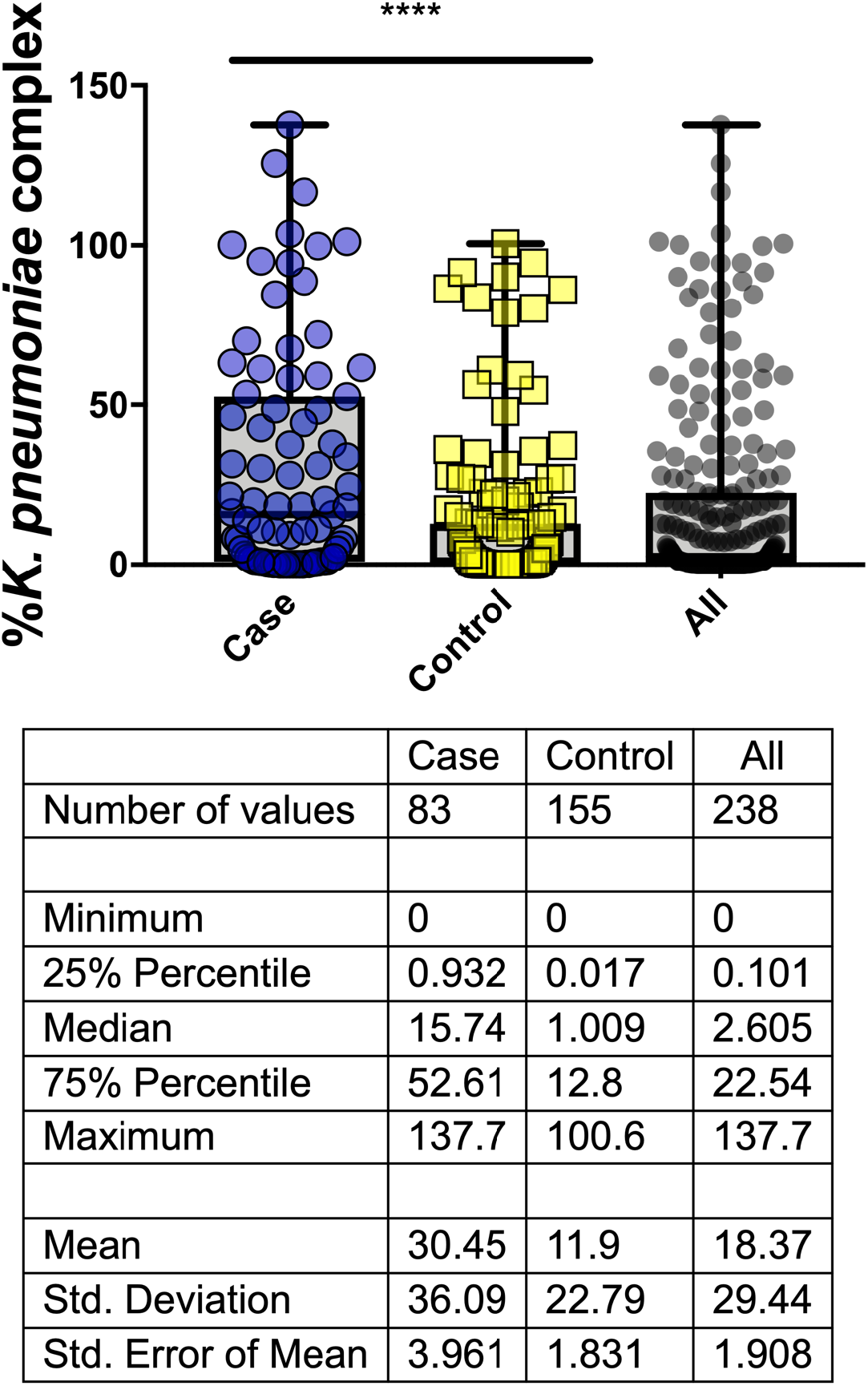
Increased relative abundance of *Klebsiella* is associated with subsequent infection. The relative abundance of *Klebsiella* in rectal swabs as measured by the Kp qPCR assay is shown for 238 specimens, with 83 cases matched 1:2 to 155 controls based on age, sex and date of swab collection. Median and interquartile ranges are shown. **** P <0.0001 by unadjusted conditional logistic regression.

### Precision and Accuracy

To assess the precision and accuracy of Kp qPCR assay, a mixture of 89% *Bacteroides ovatus*, 10% KPPR1, and 1% *Serratia marcescens* by DNA quantifications was made and diluted 10-fold serially. The mixture was amplified by the Kp qPCR assay and the relative abundances of *Klebsiella* were calculated relative to KPPR1 using the ddCT method. (**Figure 4**) The relative abundances of *Klebsiella* ranged from 15.0-18.1%, with a mean value of 17.0% (expect 10.0%). The standard deviations (hereafter SD) ranged from 0.246 to 7.386. The quantifications by Kp qPCR assay were consistent across concentrations of 4 log_10_ differences. When the total DNA concentrations of the mixture were over 1 × 10^−2^ ng/μl, the SDs of the relative abundances of *Klebsiella* were less than 1.5%. However, at 1 × 10^−3^ ng/μl total DNA concentration, the assay became less precise, as the SD increased to 7.39%. At 1 × 10^−4^ ng/μl total DNA concentration, *ybiL* assay did not detect *Klebsiella*. The copy number of 23S rRNA gene is organism-specific with 5, 7, and 8 copies in *Bacteroides ovatus*, *Serratia marcescens* and *K. pneumoniae* respectively. After adjustment for these differences, the calculated relative abundance of *Klebsiella* ranged from 10.0-12.0%, with a mean value of 11.3% (SD 0.163 to 4.91), demonstrating that Kp qPCR assay can accurately quantify *Klebsiella* from contrived samples. Fortunately, a bias in the calculation based on 23S copy number in the overall population relative to *Klebsiella* would not be expected to impact the ability to measure relative differences, as demonstrated below.

### Accuracy: Relative differences

To assess Kp qPCR assay’s ability to distinguish different relative abundances of *Klebsiella*, KPPR1 and *Escherichia coli* O6:K2:H1 CFT073 were mixed in Amies media according to CFU counts to make 10%, 20%, 30% and 40% *Klebsiella* mixtures. Ten-fold serial dilutions were made from each mixture and genomic DNA of each serial dilution were isolated and then amplified by Kp qPCR assay. The relative abundance of *Klebsiella* was calculated relative to KPPR1 using the ddCT method. (**Figure 5**) At total DNA concentrations from ~0.02-2 ng/μL the assay was able to detect relative differences between all dilutions, and at .002 ng/μL it can tell all differences except between 30 and 40% *Klebsiella.* At all but the lowest total bacterial concentration, the assay can reliably detect 10% differences in *Klebsiella* relative abundance.

### Accuracy: In Comparison to 16S rRNA sequencing

To compare the relative abundance calculated by qPCR to the gold standard of 16S rRNA sequencing, 26 samples with 16S rRNA sequence data were analyzed (**Figure 6**). *Klebsiella* relative abundances by Kp qPCR were highly correlated with 16S rRNA sequencing (Spearman’s ρ=0.79; P <.001). However, 16S sequencing does not have the resolution to separate all species and as a result, the OTU that contained *K. pneumoniae* complex also included other *Klebsiella* spp. as well as sequences from other genera such as *Enterobacter* spp. This might explain why in some cases *Klebsiella* relative abundances by Kp qPCR were significantly lower than that by 16S sequencing.

### Case-control study

To assess the association between *Klebsiella* colonization density and subsequent infection, a nested case-control study was performed. We previously enrolled 1978 subjects from 2087 separate admissions with *Klebsiella* colonization in a rectal swab, collected as part of routine surveillance for vancomycin-resistant *Enterococcus* in intensive care and oncology wards (Companion manuscript). Of these colonized subjects, 83 cases were identified that met clinical definitions of subsequent infection and had an infecting isolate that matched a colonizing isolate on or prior to the day of infection by *wzi* sequence typing. This included 41 bloodstream infections, 19 respiratory infections, and 23 urinary tract infections. Controls were defined as colonized patients who had no documented *Klebsiella* infection but had a negative clinical culture collected of the same type as the matching case. Each case was matched to two controls based on age, sex, date of rectal swab collection and swab availability, for a total of 155 controls. To find matches for every case, the criteria for age were modified for 2 cases (±20 years), and the criteria for swab collection date were modified for 4 cases (±118 days). Cases had a significantly higher comorbidity score, and were more likely to have exposure to diuretics, vitamin D, a vasopressin blocker prior to the rectal swab. They were also more likely to have been exposed to high-risk antibiotics associated with disruption of the intestinal microbiome (10). Cases also had lower baseline albumin, and were more likely to have a urinary catheter or feeding tube prior to the rectal swab.

To assess colonization density, the Kp qPCR assay was performed on all of the rectal swab samples (Figure 7). The median relative abundance was 0.022 overall, with an interquartile range (IQ) of 0.001-0.22 and an overall range of 0 to 1.38. In cases, the median was 0.15 (IQ 0.009-0.53) and in controls the median was 0.01 (IQ 0.0002-0.13). To determine if dominance was associated with infection, while accounting for the case-control matching, a cutoff for dominance was applied. The 75^th^ percentile of 0.22 in the overall dataset was chosen, consistent with cutoffs of 0.22 and 0.3 used in previous studies (6, 8). Subjects with a *K. pneumoniae* gut colonization density >22% had a 3.34-fold (1.95-5.72, P <0.0001) increased odds of infection compared to those with lower colonization density levels.

To adjust for patient variables associated with *Klebsiella* infection, inverse probability of treatment weighting was used. In the overall cohort of 1978 subjects, an explanatory model for invasive infection was built using baseline clinical features at the time of colonization (Companion manuscript, Supplemental Table 2). The model built by purposeful selection selected the following variables for inclusion: Elixhauser score, depression, diuretic use, vitamin D use, use of pressors/inotropes, low serum albumin (<2.5 g/dL), and exposure to antibiotics with high risk of microbiome disruption. These variables were then used to model *K. pneumoniae* gut colonization density >22% to generate weights for the nested case-control study. Using weights derived from these clinical covariates (available in 228 of 240 subjects), patients with a *K. pneumoniae* gut colonization density >22% had a 2.87-fold (1.64-5.03, P =0.0003) increased odds of infection compared to those with lower colonization density levels. In a secondary analysis by site of infection, increased relative abundance was also significantly associated with bloodstream infection (OR 4.137, 95% CI 1.448-11.818, *P*=0.0084), whereas associations with urinary tract (OR 3.037, 95% CI 0.571-16.17, P=0.19) and respiratory infections (OR 1.32, 95% CI 0.38-4.565, P=0.66) did not reach significance.

## Discussion

The goal of this study was to measure the association between *Klebsiella* colonization density and subsequent infection. To develop a robust method that accurately and precisely measured the relative abundance of *K. pneumoniae*, *K. quasipneumoniae* and *K. variicola* among the gut microbiota, we developed a novel qPCR assay for detecting these dominant members of the *K. pneumoniae* complex and combined it with measurement of 23S rRNA gene copies. Analytical validation indicated that this assay is inclusive of multiple strains of each species and is able to distinguish as little as 10% differences in relative abundance between samples. Applying this assay to a case-control study of *Klebsiella* infections among colonized, intensive care patients indicated that increased *Klebsiella* density is associated with subsequent infection in both unadjusted and adjusted analysis.

The finding that *Klebsiella* colonization density is associated with subsequent infection raises several interesting possibilities. One is that infection risk is dictated by how much *Klebsiella* is present in the gut, independent of the varying gene content of *Klebsiella* strains. Indeed, we and others have shown that detectable colonization is associated with infection (4, 5), and the lower limit of detection for culture is an indirect measure of abundance in rectal swabs. However, we have also demonstrated that particular *Klebsiella* genes are associated with infection as opposed to asymptomatic colonization (11), indicating that which strain a patient is colonized with affects their risk. There is likely to be an important interaction between *Klebsiella* gene content and colonization density, where certain genes may increase gut fitness and therefore gut abundance. Alternatively, there may be strains where gut abundance is increased based on microbiome factors extrinsic to *Klebsiella* but the risk of infection is further increased by virulence genes that act at the site of infection. Finally, *Klebsiella* strains with fitness genes that increase abundance in the gut and virulence genes that act later in pathogenesis are likely to pose the greatest risk of infection in colonized patients.

The main limitation of this study was the relatively small number of cases (n=83). We compensated for this by using a case-control design and an inverse probability of treatment weighting to account for clinical variables potentially associated with infection and intestinal dominance without significant loss of statistical power (12). However, we were limited in our ability to investigate associations by site of infection. Bloodstream infections were the largest infection type and were independently associated with intestinal dominance. Intriguingly, the point estimate for urinary tract infections (3.037) was similar to the overall odds ratio (2.87) for infection. This may indicate that intestinal dominance is also associated with UTIs, perhaps because a key step in pathogenesis is thought to be transit of intestinal bacteria across the perineum to the urethra.

This study further supports the growing paradigm that intestinal dominance can be used to predict infections in our hospitals. Previous studies have demonstrated that dominance of carbapenemase-producing *K. pneumoniae* in long-term care patients (8), and *K. pneumoniae* in allogeneic stem cell transplant patients (13) are associated with infection. This study evaluated a more heterogeneous population of intensive care patients with a combined outcome of bloodstream, respiratory or urinary tract infections and found the same association. The successful use of qPCR demonstrates the feasibility of measuring relative abundance of targeted pathogens in the gut using methods that are standard in clinical microbiology laboratories and inexpensive relative to next-generation sequencing. This a key step in moving towards infection prevention in hospitalized patients. A qPCR could be applied to detect colonization in rectal swabs as well as quantify it in a single step, thereby incorporating two levels of *Klebsiella* infection risk. Combined with the assessment of patient risk factors and perhaps targeted testing for *Klebsiella* virulence genes, an integrated risk assessment could be performed. If this relative risk is high enough, infection prevention interventions should be considered. Fortunately, safe and effective therapeutic strategies to eliminate gut colonization by pathogens are emerging and results from fecal transplant studies are encouraging (14). In the near future, it may be possible to assess the risk of a carbapenem-resistant *Klebsiella* infection at the time of hospital admission and prevent it without the use of antibiotics.

## Methods

### Study design and subject enrollment

This study was approved by the Institutional Review Boards of the University of Michigan Medical School (IRBMED). To assess the role of *Klebsiella* colonization density on risk of subsequent invasive infection, we conducted a nested case-control study drawn from a larger cohort of 1978 patients consecutively enrolled from 2087 inpatient admissions. Subjects admitted to intensive care units and oncology wards at our hospital undergo routine surveillance by rectal swab culture for vancomycin-resistant *Enterococcus*. After such testing, we collected residual media from these swabs and enrolled subjects into our study if colonization with *K. pneumoniae* or *K. variicola* was detected by selective culture on MacConkey agar and confirmed by MALDI-TOF identification (Bruker MALDI Biotyper). Cases were identified from this larger cohort and matched to controls as described below.

### Case definitions

Michigan Medicine patients from intensive care units (ICU) and select wards (hematology, oncology, and hematopoietic stem cell transplant) with *Klebsiella* colonization based on a rectal swab culture and a positive *Klebsiella* blood, respiratory, or urine cultures were identified as putative cases. Manual chart review was conducted by the study team to decide if they met clinical definitions of pneumonia or urinary tract infections (15–19). All patients with a *Klebsiella* blood culture were considered to have an infection. For those meeting clinical case definitions of infection, the clinical isolate and preceding rectal swab isolates were evaluated for concordance by *wzi* gene sequencing as previously described (4, 20). We have previously demonstrated that *wzi* sequencing has similar discriminatory power to 7-gene multi-locus sequence typing (4). Patients with concordant infection and colonizing isolates were considered a case. Controls were defined as colonized patients who had no documented *Klebsiella* infection but had a negative culture collected of the same type as the matching case. Cases and controls were matched 1:2 based on age ±10 years, sex, and rectal swab collection date ±90 days. This study was approved by the University of Michigan Institutional Review Board.

### Samples for PCR analysis

Rectal swabs were collected using the ESwab™ Collection and Transport System (Becton Dickinson, Franklin Lakes, NJ, USA), which elutes the sample into 1mL Amies medium. Unless specified otherwise, contrived samples that were used in PCR analysis were eluted in the Amies medium as well. The 89% *Bacteroides ovatus*, 10% KPPR1, and 1% *Serratia marcescens* mixtures were suspended in ddH_2_O.

### Bacterial DNA extraction

Genomic DNA was isolated using the MagAttract PowerMicrobiome DNA/RNA kit (Qiagen, Germantown, MD, USA) on the epMotion ^®^ 5075 liquid handler (Eppendorf, Hauppauge, NY, USA). A volume of 100 uL was added into the bead plate for each rectal swab and contrived sample. Subsequent steps of DNA extraction were conducted following the manufacturer’s instructions. *Bacteroides ovatus*, KPPR1, and *Serratia marcescens* cultures were extracted using DNeasy Blood & Tissue Kits (Qiagen, Germantown, MD, USA) following the manufacturer’s instruction for Gram-negative bacteria.

### *In silico* assay design

PanelPlex™ (DNA Software, Ann Arbor, MI, USA) *in silico* analysis was performed. Thirty-One *K. pneumoniae*, *K. quasipeumoniae*, and *K. variicola* genomes were selected as “inclusivity”, 8 *K. oxytoca* and *Raoultella* strains as “near-neighbor exclusivity”, and the human genome and common members of the gut microbiome as background sequences that should not be detected by the assay (**Supplemental Dataset 1**). PanelPlex™ utilizes ThermoBLAST™ technology to scan for thermodynamically stable off-target hybridizations that cause false-positive tests.

### Quantitative PCR assay

Real-time PCR was performed using primers (Integrated DNA Technologies, Coralville, IA, USA) and probes (Thermo Fisher Scientific, Waltham, MA, USA) with sequences and concentrations listed in Table 1 in combination with QuantiTect Multiplex PCR Kit (Qiagen, Germantown, MD, USA). PMAxx (Biotium, Fremont, CA, USA) with a final concentration of 6uM was added. A volume of 5 μL was used for each template. The final reaction volume was 25 μL. Prior to template addition, the reaction mixture was incubated for 10 minutes at room temperature and then treated in a Biotium PMA-Lite LED photolysis device for 10 minutes. PCR conditions were 50°C for 2 minutes, 95°C for 15 minutes, then 40 cycles of 94°C for 1 minute and 60°C for 1 minute on a QuantStudio 3 real-time thermocycler (Thermo Fisher Scientific, Waltham, MA, USA). KPPR1 genomic DNA was used as positive control and 100% reference for calculating *Klebsiella* relative abundances. Relative abundance was calculated using the ddC_T_ method, i.e. 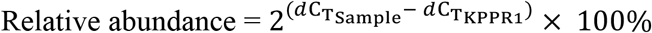, where 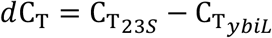.

### Statistical Analysis

Linearity was validated by linear regression. Spearman’s rank correlation coefficient was used for correlation between Kp qPCR and 16S rRNA sequencing. One-way ANOVA and Tukey’s post test was performed to compare each dilution of the KPPR1 and *Escherichia coli* O6:K2:H1 CFT073 mixtures. Statistical analysis was performed by Prism 8 (GraphPad, San Diego, CA, USA).

### Clinical modeling

Conditional logistic regression was used to study the effect of relative abundance of colonization on *Klebsiella* infection, while accounting for the case-control matching. Unadjusted analysis was performed after dichotomizing the relative abundance at the third quartile of 22%. To adjust for patient variables associated with *Klebsiella* infection, an inverse probability of treatment weighting approach was used. However, given the smaller sample size in our nested case-control study, we turned to the larger cohort from which our subjects were derived to identify most robustly the clinical variables that best explain risk of infection (Companion manuscript). First, using the increased power afforded by the overall cohort of 1978 subjects, an explanatory unconditional logistic regression model for invasive infection was built using baseline clinical features at the time of colonization. The model was built by purposeful selection, a common technique(21). Briefly, purposeful selection begins with an unadjusted analysis of each variable to select candidates with statistically significant associations with the outcome, and these are included in the starting set of covariates for the multivariable model. Iteratively, covariates are then removed from the model if they are non-significant (*P* >.05) and not a confounder (i.e. do not affect the estimate of other variables’ coefficients by at least 20%). A change in a parameter estimate above the specified level indicates that the excluded variable was important in the sense of providing a needed adjustment for one or more of the variables remaining in the model (i.e. it should be retained even if not significant). The resulting model contains significant covariates and other confounders, and then variables not included are added back one at a time. Once again the model is iteratively reduced as before but only for the variables that were additionally added. At the end of this final step, we are left with a multivariable model for *Klebsiella* infection drawn from the larger cohort of subjects with rectal *Klebsiella* colonization. The variables selected for inclusion by this method were then used to generate propensity scores for *Klebsiella* colonization density >22%, but only for subjects in the nested case-control study, again via unconditional logistic regression. The propensity scores were then used to generate weights for the IPTW process and subsequent weighted conditional logistic regression for *Klebsiella* infection The inverse of these propensity scores were then used as weights in the subsequent weighted conditional logistic regression for *Klebsiella* infection with robust standard errors. Both unadjusted and adjusted analysis were conducted using *proc survveylogistic* procedure in SAS (version 9.4), and covariate balance was assessed using the *cobalt* package in R.

## Data Availability

16S sequencing samples from rectal swabs PR08714, PR11216, PR05497, PR09929, PR10907, PR05713, PR06107, PR08411, PR08133, PR05629, PR08147, PR07331, PR12066, PR07876, PR08427, PR07976, PR08661, PR05017, PR08962, PR09113, PR08102, PR09612, PR08748, PR08048, PR06316, PR10214 are available in the sequence read archive (<u>PRJNA641414</u>).

Deidentified data from human subjects may be made available upon request, pending approval from the University of Michigan Institutional Review Board.

## Acknowledgements

MAB would like to thank Caitlyn Holmes for assistance with figure formatting.

Research reported in this publication was supported by the National Institute Of Allergy And Infectious Diseases of the National Institutes of Health under Award Number R01AI125307. The content is solely the responsibility of the authors and does not necessarily represent the official views of the National Institutes of Health.

